# *NF1* deficiency drives metabolic reprogramming in ER+ breast cancer

**DOI:** 10.1101/2023.11.24.568339

**Authors:** Rachel (Rae) J House, Elizabeth A. Tovar, Curt J. Essenburg, Patrick S. Dischinger, Abigail E. Ellis, Ian Beddows, Ryan D. Sheldon, Evan C. Lien, Carrie R. Graveel, Matthew R. Steensma

## Abstract

**Objective:** *NF1* is a tumor suppressor gene and its protein product, neurofibromin, is the key negative regulator of the RAS pathway. *NF1* is one of the top driver mutations in sporadic breast cancer such that 27% of breast cancers exhibit damaging *NF1* alterations. *NF1* loss-of-function is a frequent event in the genomic evolution of estrogen receptor (ER)+ breast cancer metastasis and endocrine resistance.

Individuals with Neurofibromatosis type 1 (NF) – a disorder caused by germline *NF1* mutations – have an increased risk of dying from breast cancer [1–4]. NF-related breast cancers are associated with decreased overall survival compared to sporadic breast cancer. Despite numerous studies interrogating the role of RAS mutations in tumor metabolism, no study has comprehensively profiled the *NF1*-mutant breast cancer metabolome to define patterns of energetic and metabolic reprogramming. The goals of this investigation were (1) to define the role of *NF1* deficiency in estrogen receptor-positive (ER+) breast cancer metabolic reprogramming and (2) to identify potential targeted pathway and metabolic inhibitor combination therapies for *NF1*-deficient ER+ breast cancer.

**Methods:** We employed two ER+ *NF1*-deficient breast cancer models: (1) an *NF1*-mutant MCF7 breast cancer cell line to model sporadic breast cancer, and (2) three distinct, *Nf1*-deficient rat models to model NF-related breast cancer [1]. IncuCyte proliferation analysis was used to measure the effect of *NF1* deficiency on cell proliferation and drug response. Protein quantity was assessed by Western Blot analysis. We then used RNAseq to investigate the transcriptional effect of *NF1* deficiency on global and metabolism-related transcription. We measured cellular energetics using Agilent Seahorse XF-96 Glyco Stress Test and Mito Stress Test assays. We performed stable isotope labeling and measured [U-^13^C]- glucose and [U-^13^C]-glutamine metabolite incorporation and measured total metabolite pools using mass spectrometry. Lastly, we used a Bliss synergy model to investigate *NF1*-driven changes in targeted and metabolic inhibitor synergy.

**Results:** Our results revealed that *NF1* deficiency enhanced cell proliferation, altered neurofibromin expression, and increased RAS and PI3K/AKT pathway signaling while constraining oxidative ATP production and restricting energetic flexibility. Neurofibromin deficiency also increased glutamine influx into TCA intermediates and dramatically increased lipid pools, especially triglycerides (TG). Lastly, *NF1* deficiency alters the synergy between metabolic inhibitors and traditional targeted inhibitors. This includes increased synergy with inhibitors targeting glycolysis, glutamine metabolism, mitochondrial fatty acid transport, and TG synthesis.

**Conclusions:** *NF1* deficiency drives metabolic reprogramming in ER+ breast cancer. This reprogramming is characterized by oxidative ATP constraints, glutamine TCA influx, and lipid pool expansion, and these metabolic changes introduce novel metabolic-to-targeted inhibitor synergies.

**HIGHLIGHTS:**

- *NF1* deficiency drives metabolic reprogramming in ER+ breast cancer.
- *NF1*-driven metabolic reprogramming is characterized by oxidative ATP constraints, glutamine TCA influx, and lipid pool expansion.
- *NF1*-deficient ER+ breast cancer cells have increased sensitivity to a combination of RAS and triglyceride synthesis inhibitors.

**Graphical Abstract:** 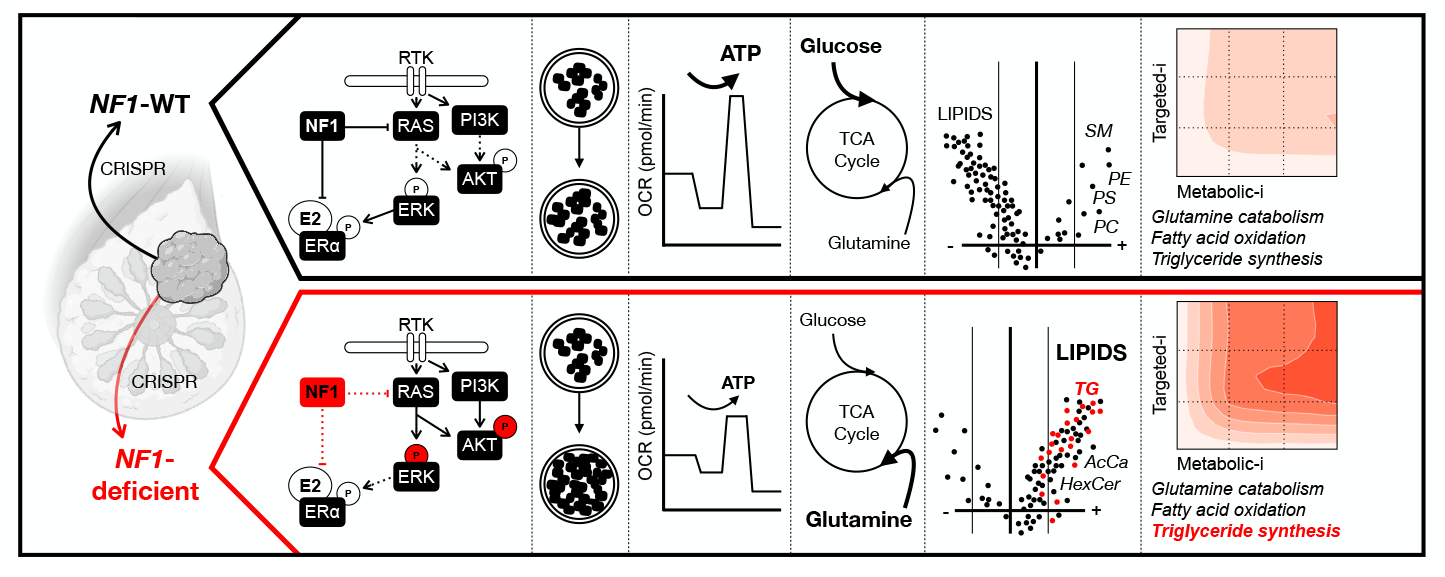

## INTRODUCTION

Recent genomic analyses have identified that the tumor suppressor gene neurofibromin 1 *(NF1)* is a driver of inherited breast cancer, sporadic breast cancer, and endocrine resistance [2–10]. The *NF1* gene product, neurofibromin, is the primary human rat sarcoma viral oncogene homolog (RAS) GTPase activating protein (GAP) and negatively regulates RAS pathway signaling. Neurofibromin’s GAP-related domain (GRD) binds RAS and stimulates RAS GTPase activity [1; 11]. *NF1* loss-of-function mutations and genomic alterations cause “*NF1* deficiency” and lead to RAS pathway dysregulation [11; 12].

Germline *NF1* mutations cause Neurofibromatosis Type 1 (NF), the most common autosomal dominant single-gene disorder, affecting 1:3000 live births [11]. Both men and women with NF have an increased risk of dying from breast cancer [13–16]. Women <40 years old with NF have a 6.5-fold increased risk of breast cancer and a 2-fold higher lifetime risk of breast cancer compared to the general population [15]. NF-related breast cancer is associated with adverse prognostic factors and decreased overall survival compared to sporadic breast cancer [13; 14; 17]. Recent studies have revealed that *NF1* is a genetic driver of sporadic breast cancer [8–10], and *NF1* shallow deletions are present in up to 27% of breast cancers [1]. Moreso, *NF1* loss-of-function mutation is a frequent event in the evolution of endocrine-resistant estrogen receptor-positive (ER+) breast cancer [4; 18]. RAS signaling is hyperactivated in approximately 50% of tumors, but RAS mutations are only found in 4% of breast cancers, suggesting that *NF1* loss of function may be driving many RAS-active breast cancers [19; 20]. These results emphasize that *NF1* deficiency and deregulated RAS signaling are critical events in breast cancer and in resistance to endocrine therapy.

Activated oncogenes and inactivated tumor suppressors contribute to tumorigenesis by reprogramming cellular energetics and metabolism towards macromolecular synthesis – a phenomenon termed “metabolic reprogramming” [21; 22]. A canonical example of cancer metabolic reprogramming is the Warburg Effect, defined as a shift towards aerobic glycolysis to support biosynthetic processes [21; 23; 24]. Metabolic reprogramming can introduce novel therapeutic targets [21; 25-32]. It is well established that RAS-activating mutations drive metabolic programming and introduce energetic and metabolic dependencies that can be therapeutically exploited [25–32]. For example, *RAS*-mutant cells are vulnerable to glutamine depletion and autophagy inhibition [30; 33-35]. Targeting RAS-driven metabolic reprogramming can also sensitize cells to targeted pathway inhibitors, demonstrating therapeutic synergy between metabolic and non-metabolic inhibitors [28; 30].

Despite numerous studies interrogating the role of RAS mutation in tumor metabolism, no study has investigated *NF1*-driven metabolic reprogramming in breast cancer. Of the limited number of *NF1*- related metabolic analyses, most have been done in *NF1* knock-out (KO) systems. Although valuable, KO models don’t account for RASGAP-independent effects. Neurofibromin’s GRD accounts for only 10% of the total neurofibromin protein, but many studies assume functional equivalence between *NF1* KO and GRD inactivation. Additionally, mutations throughout the *NF1* gene are NF and cancer-causal despite differential RASGAP effect. This data taken together underscores the utility of *NF1* mutational analyses. *NF1-*KO models have been shown to decrease mitochondrial respiration [36; 37], increase nutrient scavenging [38; 39], increase lipid droplets [40], increase resting energy expenditure [41], and induce a switch from carbohydrate to lipid catabolism [40; 41]. In non-breast cancer models, *NF1*- mutation has been shown to increase glutamine [42] and β-oxidation dependence [43]. Additionally, individuals affected by NF have lower fasting blood glucose, lower body mass indexes, increased lipid oxidation, and decreased carbohydrate catabolism compared to controls [44–46]. These analyses underscore the necessity of defining and targeting *NF1*-driven metabolic reprogramming in breast cancer.

The goals of this investigation were (1) to define the role of *NF1* deficiency in ER+ breast cancer metabolic reprogramming and (2) to identify potential targeted pathway and metabolic inhibitor combination therapies for *NF1*-deficient ER+ breast cancer. Our results revealed that *NF1* deficiency enhanced cell proliferation, altered neurofibromin isoform expression, and increased RAS and PI3K/AKT pathway signaling while constraining oxidative adenosine triphosphate (ATP) production and restricting energetic flexibility. *NF1* deficiency also increased glutamine influx into TCA intermediates and dramatically increased lipid pools, particularly triglycerides (TG). Lastly, we demonstrate that *NF1* deficiency is associated with a synergistic response to a combination of RAS pathway and TG synthesis inhibitors in clinically relevant breast cancer models.

## METHODS

### 2.1 Generation of an *NF1* sporadic human breast cancer model

To evaluate the energetic and metabolic effect of *NF1* deficiency in ER+ sporadic human breast cancer, we utilized the ER+ MCF7 human adenocarcinoma cell line. We targeted *NF1*’s RASGAP GRD domain with clustered regularly interspaced short palindromic repeat (CRISPR) guides to the 21^st^ exon and established multiple *NF1*-deficient cell lines of which one was chosen for this analysis, NF1-45 (Figure 1A). The NF1-45 mutant cells have a 5-bp deletion that introduces a premature stop codon, resulting in a truncated protein product after exon 21 (Figure 1A).

**Figure 1:**
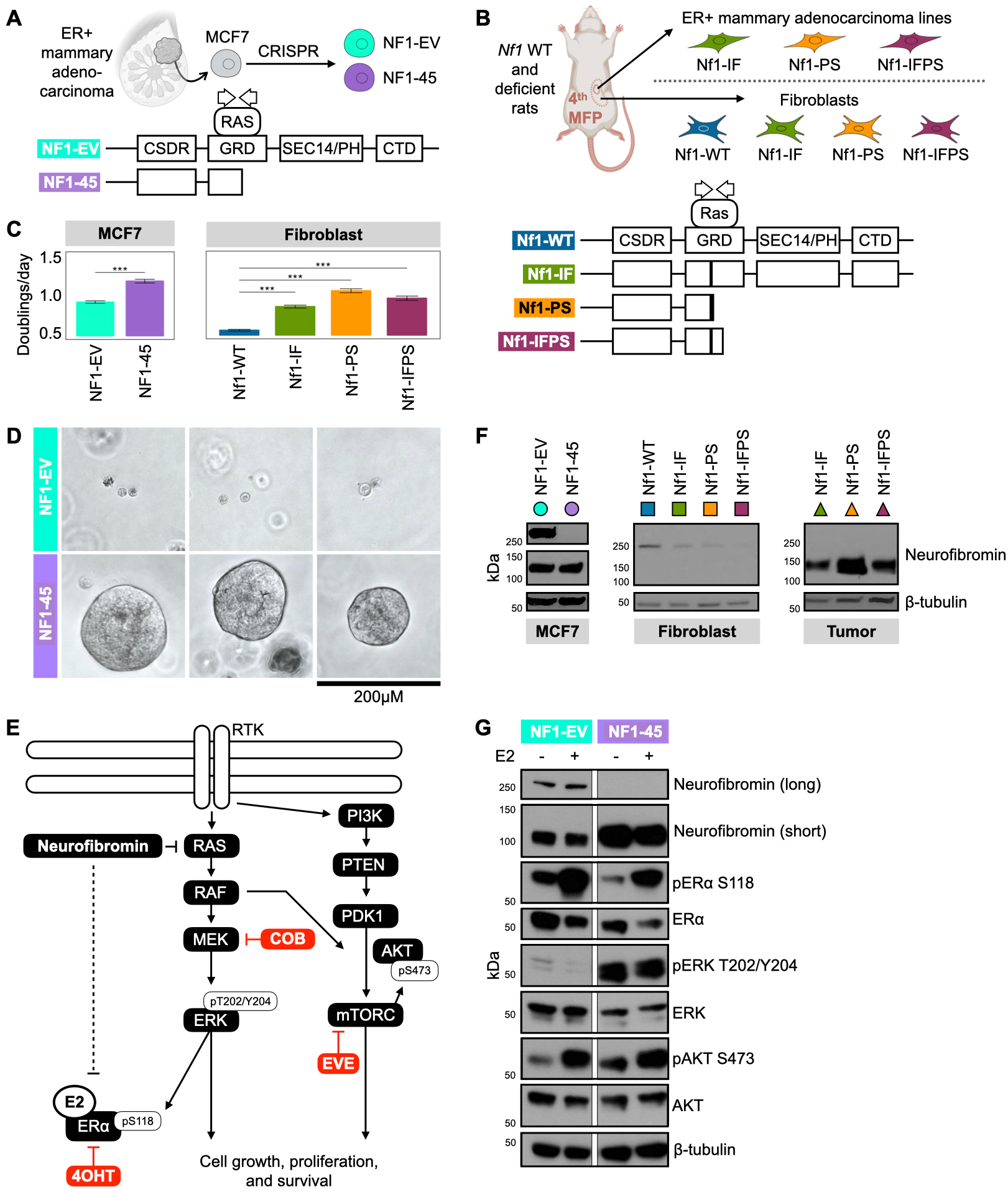
*NF1* deficiency increases cell proliferation and increases RAS and PI3K/AKT pathway signaling. A) Schematic of the *NF1*-deficient human breast cancer MCF7 model. Schematic of wild- type and mutant neurofibromin in the MCF7 NF1-EV and NF1-45 respectively. The G1 and G2 arrows within the GRD denote the CRISPR guide positioning. B) Schematic of *Nf1*-deficient rat and rat-derived fibroblast and tumor cell line models. Schematic of wild-type and mutant neurofibromin in the Nf1-WT, Nf1-IF, Nf1-PS, Nf1-IFPS rat models. The G1 and G2 arrows within the GRD denote the CRISPR guide positioning. C) *NF1* deficiency increases cell proliferation in MCF7 cells and rat mammary fibroblasts, n=2, 24 tech reps per n. The bar represents the geometric mean, and the black, horizontal lines represent the 95% confidence interval boundaries. D) *NF1*-deficient MUT-45 spheroids grow larger than NF1-EV spheroids when grown in Matrigel. (n=3+, 1 representative technical replicate shown above, day 10 of 3D culture). E) Schematic description of the role of neurofibromin in the ER, RAS, and PI3K/AKT signaling pathways and relevant targeted inhibitors: Cobimetinib (COB), Tamoxifen (4OHT), and Everolimus (EVE). F) *NF1* indels alter neurofibromin expression. Western blot analysis of total neurofibromin protein. Immunoblotting was carried out using antibodies against neurofibromin (H12, Bethyl, iNF-07E) and β-tubulin (n=3+, 1 representative biological replicate shown). G) *NF1* deficiency increases RAS and AKT/PI3K pathway signaling. Western blot analysis of ER, RAS, and PI3K/AKT phospho-signaling. Immunoblotting was carried out using antibodies against neurofibromin (Bethyl), pERα S118, ERα, pERK T202/Y204, ERK, pAKT S473, AKT, and β-tubulin (n=3, 1 representative biological replicate shown).

### 2.2 NF-related rat breast cancer models

To understand the energetic and metabolic effect of *Nf1* deficiency in NF-related breast cancer, we employed our previously developed and described NF-related spontaneous rat mammary adenocarcinoma models [1]. To generate these models, we targeted *Nf1*’s RASGAP GRD domain with CRISPR guides to the 20^th^ exon. We generated three distinct *Nf1* indels in exon 20 that caused either in-frame or premature stop deletions. The *Nf1* inframe mutant (Nf1- IF) rats have a 54bp deletion in exon 20 (Figure 1B) [1]. The *Nf1* premature stop mutant (Nf1-PS) rats have an 8-bp deletion in exon 20 that introduces a premature stop codon, resulting in a truncated protein product after exon 20 (Figure 1B) [1]. The *Nf1* inframe premature stop mutant (Nf1-IFPS) rats have a 57-bp deletion in exon 20 that results in a unique messenger RNA (mRNA) isoform with an additional 140-bp deletion in exon 21 [1]. The exon n21 mRNA deletion introduces a premature stop codon, resulting in a truncated protein product after exon 21 (Figure 1B) [1]. The IF, PS, and IFPS indels model “*Nf1* deficiency” and induce aggressive, 100% penetrant, multi-focal, ER/PR+ mammary adenocarcinomas in both female and male rats [1]. To perform *in vitro* energetic and metabolic experiments using our NF-related rat breast cancer model, we utilized isolated, *ex vivo* rat mammary fat pad fibroblasts from each genetic background: Nf1-WT, Nf1-IF, Nf1-PS, Nf1-IFPS [47]. We also used isolated, *ex vivo* adenocarcinoma cell lines from each of the *Nf1*-deficient rat models: Nf1-IF, Nf1-PS, Nf1-IFPS [47]. Although mammary epithelial cells (MECs) are our proposed tumor cell of origin, our isolated MECs do not passage *in vitro*, so we used isolated fibroblasts to have a rat *ex vivo Nf1*-WT to *Nf1*-deficient control since the rat tumor cell lines do not have a *Nf1*-WT control.

### 2.3 Cell Culture

Human ER+ MCF7 breast cancer cells (HTB-22), were grown in Dulbecco’s modified eagle medium (DMEM) no phenol red (Gibco), 10% fetal bovine serum (FBS), 5mL penicillin-streptomycin (Gibco). Rat-derived fibroblast lines were grown in DMEM/F12 no phenol red (Gibco), 10% FBS, and 5mL gentamycin (Gibco). Rat adenocarcinoma-derived tumor cell lines were grown in MEC media (DMEM/F12 no phenol red, 10% horse serum, 20ng/mL EGF, 0.5ug/mL hydrocortisone, 100ng/mL cholera toxin, and 10ug/mL insulin). All cells were grown at 37°C in a 5% CO2 incubator. Mycoplasma testing (ATCC) every 6 months confirmed cells were not infected.

### 2.4 IncuCyte proliferation analysis

Cells were grown in their respective medias (see section 2.3) and treated with 10nM beta-estradiol for 48hr in a Sartorius IncuCyte. Cell proliferation was measured via cell confluency and doublings per day were calculated via the equation: Doublings per day = (final confluency / starting confluency) / assay duration (2 days).

### 2.5. 3-Dimensional Culture of Cells on Matrigel

MCF7 cells were plated into 8-well chamber slides at a density of 2×10^4^ cells/well in 100uL GFR-Matrigel. After Matrigel solidification, overlay 3D MEC media was added. Cells were fed with fresh media every 3 days and imaged on Day 3, 6, and 10. Day 10 is shown.

### 2.6 RNAseq

RNAseq was performed on MCF7 NF1-EV and NF1-45 cell lines using 3 biological replicates. For each technical replicate, 1.2×10^5^ cells were plated into a well of a 6-well plate. After 24hr. RLT buffer (RNeasy kit) was added to each well. The RLT buffer and cells were added to an MPbio lysing matrix E tube, and cells were homogenized for 20s in an MPbio homogenizer. The supernatant was transferred to a QIAshredder column, and an equal volume of 70% ethanol was added to the flow-through. The flow-through and ethanol mixture was transferred to an RNeasy column and the RNA was extracted following the manufacturer’s instructions. The Van Andel Genomics Core prepared RNA libraries from 500ng of total RNA using a KAPA mRNA Hyperprep kit (v4.17, Kapa Biosystems). RNA was sheared to 300-400bp. Before amplification, cDNA fragments were ligated to IDT for Illumina TruSeq UD Indexed adapters (Illumina Inc). Library quality was assessed using Aglient DNA High Sensitivity chip (Agilent Technologies, Inc.), QuantiFluor ® dsDNA System (Promega Corp.), and Kapa Illumina Library Quantification qPCR assays (Kapa Biosystems). Indexed libraries were combined and 100bp, paired-end sequences was performed using an Illumina NovaSeq6000 sequencer using a S4, 200-cycle sequencing kit (Illumina Inc.) to a depth of 70M reads per sample.

Illumina RTA3 was used for base-calling, and NCS output was demultiplexed and converted to FastQ with Illumina Bcl2fastq v1.9.0. Trim Galore v0.6.0 was used to trim sequencing adapters from demultiplexed FASTQ files (https://www.bioinformatics.babraham.ac.uk/projects/trim_galore/).

GENCODE was used to map FASTQ files to GRCh38 (release 33 with STAR v2.7.8a basic two-pass mode)[48]. Raw counts were extracted from reverse strand alignments, and principal component analysis was computed using iDEP 9.1. Fold change was calculated using normalized counts per million. Msigdbr [49] and clusterProfiler [50] packages were used to perform Gene Set Enrichment Analysis (GSEA) using 10,000 permutations and genes ranked by fold change. For the metabolism-related GSEA analysis, we manually curated a list of metabolism-related Kyoto Encyclopedia of Genes and Genomes (KEGG) and REACTOME pathways from a list of all KEGG and REACTOME pathways. The data was then queried for pathways with significantly increased or decreased enrichment across 5 or more comparisons (NF1-45 vs. NF1-EV, Nf1-IF fibroblasts vs. Nf1-WT fibroblasts, Nf1-PS fibroblasts vs. Nf1-WT fibroblasts, Nf1-IFPS fibroblasts vs. Nf1-WT fibroblasts, Nf1-IF tumor vs. Nf1-WT whole mammary fat pad (MFP), Nf1-PS tumor vs. Nf1-WT whole MFP, and Nf1-IFPS tumor vs. Nf1-WT MFP).

### 2.7 Total protein expression westerns

Cells were grown in their respective medias (see section 2.3) and treated with 10nM beta-estradiol for 48hr before lysate collection. Cells were washed with ice-cold 1X PBS, scraped, and lysed in Rb buffer (20 mM TrisHCL, pH 7.6; 5 mM EDTA; 150 mM NaCl; 0.5% NP-40; 50 mM NaF; 1 mM beta-glycerophosphate) supplemented with PhosSTOP and protease inhibitors. Samples were resolved by SDS-PAGE. Immunoblotting was carried out using antibodies against neurofibromin (Bethyl, H12, and iNF-07E), voltage-dependent anion channel (VDAC), and β-tubulin.

### 2.8 ER, RAS, and PI3K phospho signaling westerns

The MCF7 cells were grown in their aforementioned media (see section 2.3) and treated with 10nM beta-estradiol in DMEM no phenol red (Gibco), 2% charcoal-stripped FBS (cFBS), and 5mL penicillin-streptomycin for 24hr before lysate collection. Cells were washed with ice-cold 1X PBS, scraped, and lysed in Rb buffer supplemented with PhosSTOP and protease inhibitors. Samples were resolved by SDS-PAGE. Immunoblotting was carried out using antibodies against neurofibromin (Bethyl), pERα S118, ERα, pERK T202/Y204, ERK, pAKT S473, AKT, and β-tubulin (Cell Signaling).

### 2.9 Seahorse XF-96 energetic analysis

Cells were grown in their respective medias (see section 2.3) and treated with 10nM beta-estradiol for 48hr before plating for the Seahorse assay. Cells were trypsinized, collected, quenched, and spun down. Cells were counted with an automated BioRad cell counter and plated in poly-d-lysine-coated Seahorse XF-96 assay plates. Cells were plated with the goal of ∼80-90% final confluency, so MCF7 cells were plated at 30,000 cells per well, fibroblasts at 5,000 cells per well, and tumor cells at 15,000 cells per well. Cells were spun down and allowed to fully adhere to the plates for 4hr at 37°C in a 5% CO2 incubator. After 4hr, growth media was gently removed by pipetting, and 180μL Seahorse XF DMEM media was added to each well. Seahorse Mito Stress Test assay media contained 25mM glucose, 4mM glutamine, and 1mM pyruvate to replicate DMEM. Seahorse Glyco Stress Test assay media contained 4mM glutamine. The assay was performed according to Agilent Mito Stress Test and Glyco Stress Test instructions. Extracellular acidification rates (ECAR) and oxygen consumption rates (OCR) shown are per 5000 cells. ATP metrics were calculated following Mookerjee et al. 2017 [51] and using the CEAS R Package [52].

### 2.10 Stable isotope labeling and metabolomics

For stable isotope labeling experiments with *in vitro* MCF7 NF1-EV and NF1-45 cells, cells were grown in DMEM no phenol red (Gibco), 10% FBS, 5mL penicillin-streptomycin (Gibco) and treated with 10nM beta-estradiol for 48hrs. Cells were washed in PBS and incubated in DMEM no phenol red, no glucose, no glutamine (Gibco), 10% dialyzed FBS (dFBS), 5mL penicillin-streptomycin (Gibco), and 10nM beta-estradiol with either 25mM [U-^13^C]-glucose and 4mM [U-^12^C]-glutamine or 25mM [U-^12^C]-glucose and 4mM [U-^13^C]-glutamine for 2hrs. [U-^13^C]-glucose and [U-^13^C]-glutamine were purchased from Cambridge Isotope Laboratories. After 2hr, cells were washed 2X with ice-cold NaCl saline, snap-frozen on dry ice, and stored at -80°C. Metabolites were extracted using a modified Bligh Dyer method [53]. This modified Bligh Dyer extraction involves the addition of ice-cold methanol to the cell plate from which cells are scraped and transferred to an Eppendorf tube containing ice-cold chloroform and water, with a final v/v ratio of 2:2:1.8 methanol to chloroform to water. The cell extract was then vortexed for 10 seconds and incubated on wet ice for 30 minutes. The samples were then vortexed again for 10 seconds and spun down at 4°C for 10 minutes at 13,000rpm to achieve aqueous and organic phase separation. The aqueous and organic phases were collected and dried down in a speedvac.

### 2.11 LC-MS method (aqueous)

For liquid chromatography-mass spectrometry (LC-MS) analysis, the aqueous dry-down was resuspended in 50μL of 50:50 acetonitrile and water (A955, Fisher Scientific). The resuspended metabolite extracts were analyzed on an Orbitrap ID-X mass spectrometer (Thermo Fisher Scientific) with a Thermo Vanquish Horizon LC system. Each sample injection was 2μL. The Van Andel Institute Mass Spectrometry Core performed chromatographic separation using Acquity BEH Amide (1.7μm, 2.1mm x 150mm) analytical columns (#176001909, Waters, Eschborn, Germany). These columns were fitted with a pre-guard column (1.7μm, 2.1mm x 5mm; #186004799, Waters) and used an elution gradient with two solvents. Solvent A was LC/MS grade water (W6-4, Fisher), and Solvent B was 90% LC/MS grade acetonitrile (A955, Fisher). For negative mode analysis, both mobile phases contained 10 mM ammonium acetate, 0.1% ammonium hydroxide, and 5 μM medronic acid. For positive mode analysis, both mobile phases included 10 mM ammonium formate and 0.1% formic acid. The 20-minute gradient at 400 μL/min consisted of the following phases: 0–1.0 min (100% B to 90% B), 1.0–12.5 min (90% B to 75% B), 12.5–19 min (75% B to 60% B), and 19–20 min (60% B). After each run, the column was re-equilibrated over 20 minutes. The column temperature was maintained at 40°C. The heated electrospray ionization (H-ESI) source used a spray voltage of -2500 V (negative mode) or 3500 V (positive mode).

The isotopically labeled experimental replicates were collected in full-scan mode with an MS1 scan range from 70 to 1000 m/z, a mass resolution of 120,000 FWHM, RF lens at 35%, and automatic gain control. The unlabeled samples were used for data-dependent MS2 (ddMS2) fragmentation and were annotated using AquireX workflow. ddMS2 data was obtained with MS1 resolution set at 60,000, MS2 resolution at 30,000, an intensity threshold of 2.0 x 104, and dynamic exclusion activated after a single trigger for a duration of 10 seconds. MS2 fragmentation comprised HCD at 20%, 35%, and 50% collision energies, which was followed by CID at 15%, 30%, and 45% with an activation Q of 0.25. Both MS2 scans used standard automatic gain control and a maximum injection time of 54 ms. The total cycle time for MS1 and ddMS2 scans was 0.6 seconds.

### 2.12 LC-MS method (organic)

The organic dry-down was resuspended in 50μL of 50:50 isopropanol:water. The resuspended metabolite extracts were analyzed on an Orbitrap ID-X mass spectrometer (Thermo Fisher Scientific) with a Thermo Vanquish Horizon LC system. Each sample injection was 2μL. The Van Andel Institute Mass Spectrometry Core performed chromatographic separation using C30 (2.6μm, 2.1mm x 150mm) analytical columns (#27826-152130 Thermo Fisher Scientific). These columns were fitted with a pre-guard column (1.7μm, 2.1mm x 5mm; #186004799, Waters) and used an elution gradient with two solvents. Solvent A was 60% acetonitrile, 40% water, 0.1% formic acid, 10mM ammonium formate, and Solvent B was 90% isopropanol, 8% acetonitrile, 2% water, 0.1% formic acid, 10mM ammonium formate. The 30-minute gradient at 400 μL/min consisted of the following phases: 0-1.0 min (75%A), 1.0-3.0 min (75%A to 60%A), 3-19 min (60%A to 25%A), 19-20.5 min (25% A to 10% A), 20.5-28 min (10% A to 5% A), and 28.1-30 min (5% to 0%A). After each run, the column was re-equilibrated over 15 minutes. The column temperature was maintained at 50°C. The heated electrospray ionization (H-ESI) source used a spray voltage of -2800 V (negative mode) or 3250 V (positive mode). The isotopically labeled experimental replicates were collected in full-scan mode with an MS1 scan range from 200 to 1000 m/z, a mass resolution of 240,000 FWHM, RF lens at 45%, and automatic gain control. The unlabeled samples were used for data dependent MS3 (ddMS3) fragmentation. ddMS2 scans were collected with HCD fragmentation settings (Assisted Collision Energy Mode, Normalized HCD Collision Energies at 15%, 30%, 45%, 75%, and 110% energy levels, Orbitrap resolution: 15,000). A m/z 184 mass trigger, indicative of phosphatidylcholines, was used for CID fragmentation (Collision energy: 35%, activation time: 10ms, Orbitrap resolution: 15,000). CID MS3 scans were triggered by specific acyl chain losses for detailed analysis of mono-, di-, and tri- acylglycerides (Collision energy: 35%, activation time: 10ms, Ion trap detection at Rapid Scan Rate).

### 2.13 LC-MS data analysis

Lipid identifications were assigned using LipidSearch (v5.0, Thermo Fisher Scientific). Data was analyzed using Compound Discoverer (v 3.2, Thermo Fisher Scientific). Compounds were identified via retention compared external standards and MS2 spectral matching using the mzCloud database (Thermo Scientific). Mass isotopologue distribution analysis was performed using isotopologue peak area data from labeled and unlabeled samples, natural abundance correction was performed using an internal VAI matrix-based algorithm like those described in Fernandez et al.1996 and Trefely et al. 2016. PCA analysis was performed using the FactoMineR R package.

### 2.14 FBS Dialysis

To deplete glucose and glutamine from the FBS used during stable isotope labeling, we prepared dialyzed FBS. In short, FBS was pipetted into SnakeSkin™ Dialysis Tubing, 10K MWCO, 35 mm, 10.5m (Thermo # 88245). The tubing was clamped closed using SnakeSkin Dialysis clips (Thermo # 68011). The FBS-containing dialysis tubing was suspended in 1X PBS (1:300 ratio of FBS:PBS) using a foam Eppendorf tube holder. The PBS was stirred around the FBS-containing tubing using a magnetic stir bar and stir plate for 2hr at 4°C, after which the PBS was discarded and replaced. The PBS was then stirred around the FBS-containing tubing for 3hr at 4°C, after which the PBS was discarded and replaced. Lastly, the PBS was stirred around the FBS-containing tubing for 18-20hr at 4°C, after which the FBS was pipetted out of the tubing, filtered with a 0.22μM filter, aliquoted, and stored at -20°C. In total, the FBS was dialyzed against 1X PBS for approximately 24hr. Mass spectrometry was used to confirm metabolite depletion (Supplementary Fig. 2).

### 2.15 Targeted and metabolic inhibitor *in vitro* drug studies

MCF7 cells were grown in DMEM no phenol red (Gibco), 10% FBS, 5mL penicillin-streptomycin (Gibco) and treated with 10nM beta-estradiol and targeted (Tamoxifen [4OHT], Cobimetinib [COB], Everolimus [EVE]) and/or metabolic (2-Deoxy-D-glucose [2DG], bis-2-(5-phenylacetamido-1,3,4-thiadiazol-2-yl)ethyl sulfide [BPTES], Etomoxir [EX], PF-04620110, PF-06424439) inhibitors 48hr in a Sartorius IncuCyte. Cell proliferation was measured via cell confluency and doublings per day were calculated via the equation: Doublings per day = (final confluency / starting confluency) / assay duration (2 days). Synergy was calculated using the SynergyFinder R package.

## RESULTS

### 3.1 *NF1* deficiency increases cell proliferation and increases RAS and PI3K/AKT pathway signaling

To evaluate the metabolic effect of *NF1* mutation in ER+ sporadic human breast cancer, we designed and employed a novel *NF1-*deficient ER+ MCF-7 human adenocarcinoma cell line model, NF1-45 (Figure 1A). The NF1-45 mutant cells have a 5-bp GRD deletion that introduces a premature stop codon, resulting in a truncated protein product after exon 21 (Figure 1A). To evaluate the metabolic effect of *NF1* mutation in an NF context, we derived *ex vivo* fibroblast and mammary adenocarcinoma cell lines from our previously described *Nf1*-deficient rat models [1]. The *Nf1* inframe mutant (Nf1-IF) rats have a 54bp deletion in exon 20 (Figure 1B) [1]. The *Nf1* premature stop mutant (Nf1-PS) rats have an 8-bp deletion in exon 20 that introduces a premature stop codon, resulting in a truncated protein product after exon 20 (Figure 1B) [1]. The *Nf1* inframe premature stop mutant (Nf1- IFPS) rats have a 57-bp deletion in exon 20 that results in a unique messenger RNA (mRNA) isoform with an additional 140-bp deletion in exon 21 [1]. The exon n21 mRNA deletion introduces a premature stop codon, resulting in a truncated protein product after exon 21 [1]. The *Nf1* IF, PS, and IFPS indels model “*Nf1* deficiency” and induce aggressive, penetrant, multi-focal, ER/PR+ mammary adenocarcinomas in both female and male rats. The Nf1-PS rats have the earliest onset tumors at ∼8 weeks, followed by Nf1-IFPS at ∼10 weeks, followed by Nf1-IF at ∼1yr. Before performing energetic and metabolic analyses, we aimed to characterize our novel MCF7, fibroblast, and tumor cell lines. To evaluate the impact of *NF1*-deficiency on cell proliferation, cells were grown for 48hr with 10nM estradiol (E2) in a Sartorius IncuCyte, and cell doublings were measured via confluence over time. Of note, unless otherwise stated, all assays were conducted in the presence of estradiol to better replicate an *in vivo* environment. Compared to *NF1*-WT controls (NF1-EV and Nf1-WT), *NF1*-deficient cells proliferate faster in 2D (Figure 1C). When compared to the NF1-EV control, NF1-45 developed larger 3D spherical colonies, indicating greater tumor potential (Figure 1D). Since neurofibromin is the primary RAS-GAP, alteration of neurofibromin leads to dysregulated RAS-pathway signaling, and RAS pathway signaling can activate the parallel PI3K-AKT pathway (Figure 1E). Neurofibromin can also indirectly activate and directly inhibit ER signaling and alter cellular localization (Figure 1E) [54]. Given the relationship between, neurofibromin, ER, RAS, and PI3K-AKT signaling, we performed western blot analysis to characterize neurofibromin expression and ER, RAS, and PI3K-AKT signaling (Figure 1F and 1G). *NF1* deficiency resulted in decreased expression of the putative neurofibromin 250kDa isoform (Figure 1F, Supplementary Fig.1). *NF1* deficiency also increased the ratio of a novel neurofibromin 125kDa isoform to neurofibromin 250kDa isoforms in tumor lines (Figure 1F). When we measured ER signaling via phosphorylation at the ERK activation-associated S118 site, we saw that NF1-45 cells had decreased active and total ERα activation at baseline and following estradiol (E2) stimulation (Figure 1G). Conversely, when we evaluated ERK signaling, we found that *NF1* deficiency dramatically increased active, T202/Y204-phosphorylated ERK, which is indicative of increased RAS-pathway signaling (Figure 1G). We then probed against the activation-associated AKT S473 phosphorylation sites and saw a modest increase in PI3K-AKT pathway signaling (Figure 1G). To summarize our phenotyping results, *NF1* deficiency increases 2D and 3D cell proliferation, alters neurofibromin isoform expression, decreases ER signaling, and increases RAS and AKT signaling.

### 3.2 *NF1* deficiency alters global transcription and metabolism-related transcriptomic networks

To better understand the impact of *NF1* deficiency on global and metabolism-related transcription, we performed RNAseq and utilized principal component analysis (PCA) to visualize the variability across *NF1*-WT and *NF1*-deficient cells (Figure 2A). PCA displayed the global shift in gene expression in the *NF1*-deficient cells compared to the *NF1*-WT controls (Figure 2A). To evaluate the effect of *NF1* deficiency on metabolism-related transcription, we performed gene set enrichment analysis (GSEA). We queried the data for metabolism-related KEGG and REACTOME pathways that significantly increased or decreased in enrichment across our *NF1*-deficient models. We found that *NF1* deficiency altered oxidative phosphorylation-related (OXPHOS) transcription (Figure 2B), a finding that is phenotypically paralleled in our energetic analyses (Figure 3). We also found that *NF1* deficiency decreased FA metabolism, membrane lipid production-related transcription, and branched-chain amino acid degradation-related transcription (Figure 2B). Our RNAseq analysis demonstrates that *NF1* indels alter metabolism in breast cancer. Next, we sought to define the impact of *NF1* mutations on cellular energetics (glycolysis and OXPHOS) and amino acid and lipid metabolism.

**Figure 2:**
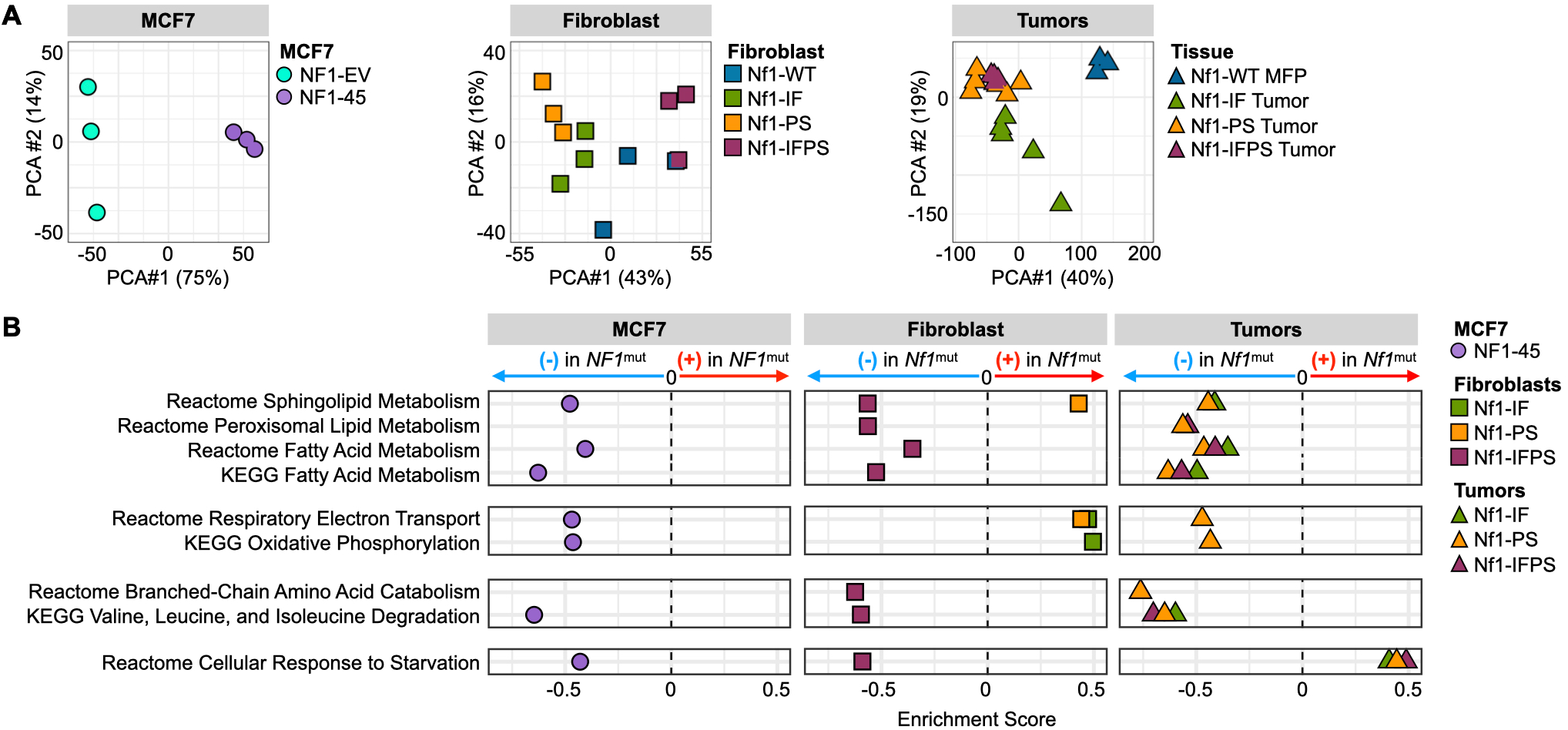
*NF1* deficiency alters global transcription and metabolism-related transcriptomic networks. A) *NF1*-deficient cells are transcriptionally distinct across cell types and models (n=3). PCA coordinates were generated using iDEP 9.1. B) *NF1* deficiency alters the metabolic transcriptome. GSEA enrichment analysis for metabolism-related KEGG and Reactome pathways. Each bubble represents the pathway enrichment score for each *NF1*-deficient vs. *NF1*-WT comparison. The list of metabolism-related KEGG and REACTOME pathways was manually curated from a list of all KEGG and REACTOME pathways in all models.

**Figure 3:**
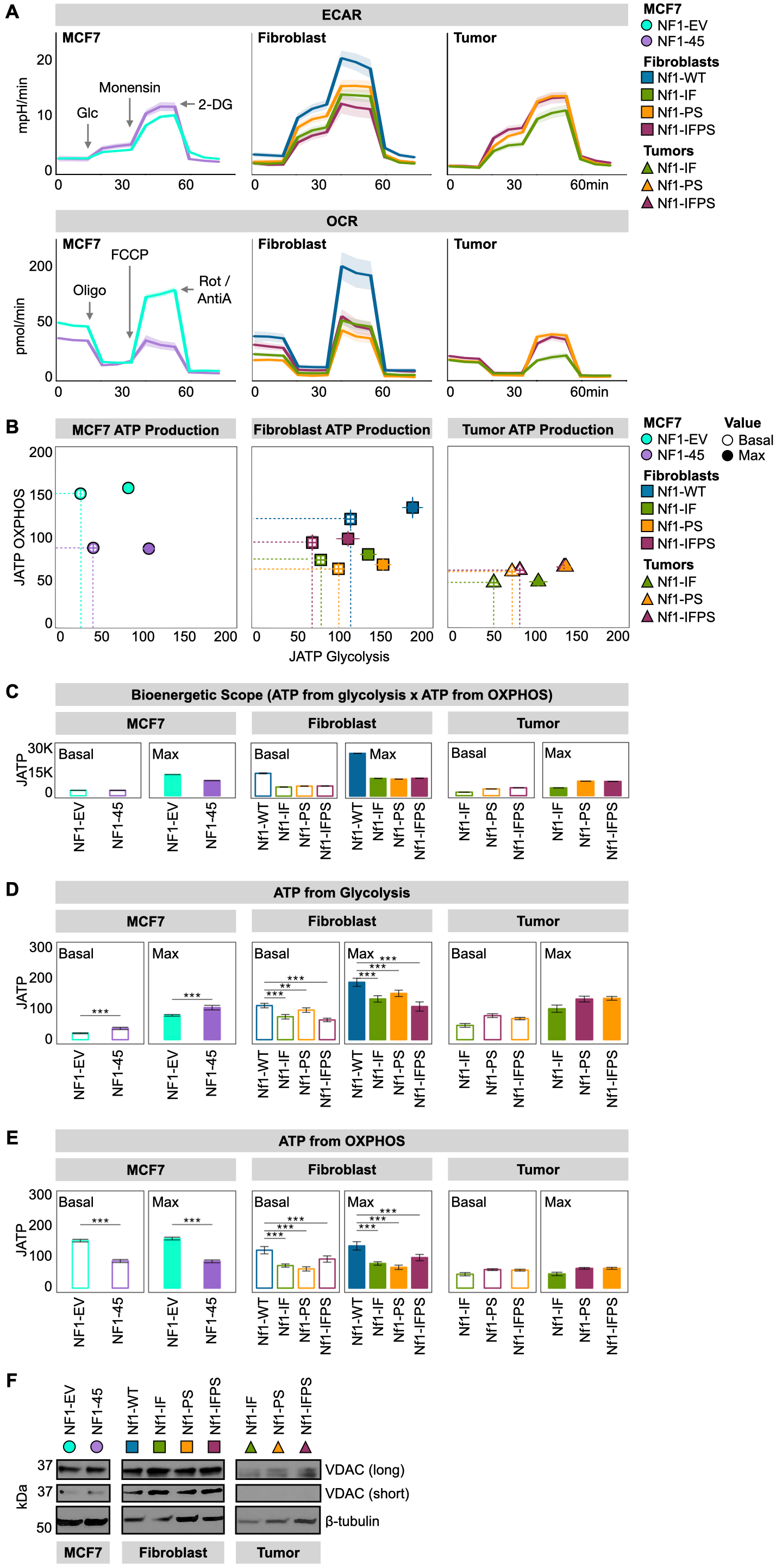
*NF1* deficiency constrains oxidative ATP production. A) *NF1* deficiency decreases basal and maximal OCR across models. Monensin is an ionophore that pushes cells to the maximal glycolytic range. 2DG is a competitive hexokinase inhibitor. Oligomycin (oligo) is a complex V inhibitor, fluoro- carbonyl cyanide phenylhydrazone (FCCP) is an uncoupling agent, and rotenone and antimycin A (Rot/AntiA) are complex I and complex III inhibitors, respectively. B) *NF1* deficiency restricts joules of ATP (JATP) production in MCF7 cells and rat mammary fibroblasts. JATP values were calculated using CEAS (R Package). Dotted lines outline the basal bioenergetic scope area. The symbols represent the geometric mean, and the lines represent the 95% confidence interval. MCF7: n = 2, 12 technical replicates per biological replicate. Fibroblast: n=2, 24 technical replicates per biological replicate. Tumor cells: n=2, 28 technical replicates per biological replicate. C) *Nf1* deficiency restricts max bioenergetic scope (JATP from glycolysis x JATP from OXPHOS) in MCF7 cells and rat mammary fibroblasts. The bar represents the geometric mean, and the black, horizontal lines represent the 95% confidence interval boundaries. JATP values were calculated using CEAS (R Package). D) *NF1* deficiency has a divergent effect on glycolytic JATP production based on cell type. JATP values were calculated using CEAS (R Package). The bar represents the geometric mean, and the black, horizontal lines represent the 95% confidence interval boundaries. MCF7: n=2, 12 technical replicates per biological replicate. Fibroblast: n=2, 24 technical replicates per biological replicate. Tumor cells: n=2, 28 technical replicates per biological replicate. E) *NF1* deficiency restricts OXPHOS JATP production. JATP values were calculated using CEAS (R Package). The bar represents the geometric mean, and the black, horizontal lines represent the 95% confidence interval boundaries. MCF7: n=2, 12 technical replicates per biological replicate. Fibroblast: n=2, 24 technical replicates per biological replicate. Tumor cells: n=2, 28 technical replicates per biological replicate. F) *NF1* deficiency impacts OCR without altering total mitochondrial load as measured by Western blot analysis of voltage-dependent anion channel (VDAC) expression. Immunoblotting was carried out using antibodies against VDAC and β-tubulin.

### 3.3 *NF1* deficiency constrains oxidative ATP production

Cellular energetics describes how energy is captured and used by cells. Our analysis focused on core aspects of energy metabolism: glycolysis, the tricarboxylic acid cycle (TCA), and the electron transport chain (ETA). To measure cellular energetics in real-time, we used the Seahorse XF-96 Mito Stress Test and Glyco Stress Test assays. The Seahorse machine has two main outputs: extracellular acidification rate (ECAR) and oxygen consumption rate (OCR). ECAR measures media acidification and is a proxy of glycolysis. OCR measures media oxygen depletion and is a proxy of OXPHOS (O2 is the final electron transport chain [ETC] electron acceptor). Cellular energetics were modulated using the following compounds: glucose (glc) is a substrate for glycolysis, monensin is an ionophore, 2-deoxy-d-glucose (2DG) is a competitive hexokinase inhibitor, oligomycin (oligo) is a complex V inhibitor, fluoro-carbonyl cyanide phenylhydrazone (FCCP) is an uncoupling agent, and rotenone and antimycin A (Rot/AntiA) are complex I and complex III inhibitors, respectively. Across our human and rat models, *NF1* deficiency decreases basal and maximal OXPHOS (Figure 3A). When we calculated the ATP produced from glycolysis and OXPHOS and visualized those values in two dimensions, we found that *NF1* deficiency restricts energetic flexibility (Figure 3B). Despite oxidative constraints, only NF1-45 cells have glycolytic “compensation” at basal conditions, but all *NF1*-deficient lines have smaller max bioenergetic scopes, a metric calculated by multiplying the JATP glycolysis x JATP OXPHOS (Figure 3C). Bioenergetic scope describes the theoretical energetic space in which a cell can operate, so the larger the bioenergetic scope, the greater the energetic flexibility. When we compared *NF1*-WT and *NF1*-deficient ATP production from glycolysis and OXPHOS, we found that *NF1*-deficient cells have decreased OXPHOS ATP production (Figure 3D and 3E). This finding underscores the OXPHOS-related transcriptomic changes seen in our RNAseq analysis (Figure 2B). To understand whether this decrease in OXPHOS capacity was due to a decrease in total mitochondrial load, we performed Western Blot analysis and analyzed voltage-dependent anion channel (VDAC) expression. When we compared our *NF1*-deficient models to their WT controls, we saw no appreciable difference in VDAC expression (Figure 3F).

### 3.4 *NF1* deficiency increases intracellular lipid pools

To define the *NF1*-deficient metabolome, we performed untargeted metabolomics and stable isotope labeling. We compared intracellular metabolite pools, labeled pool sizes (PS), and labeled pool fractional enrichment (FE) between *NF1*-WT and *NF1*-deficient MCF7 cells and visualized this difference with PCA analysis (Figure 4A). Across all queries, *NF1*-deficient cells are distinct, and this difference is most pronounced when comparing total metabolite pools (Figure 4A). Because we saw an *NF1*-driven OXPHOS loss, we wanted to understand how glucose and glutamine were incorporated into TCA intermediates via mass isotopologue distribution. [U-^13^C]-glucose TCA incorporation is facilitated by pyruvate dehydrogenase (PDH) and pyruvate carboxylase (PC), which respectively catalyze the conversion of pyruvate to acetyl-CoA and oxaloacetate. [U-^13^C]-glucose TCA entry via PDH leads to M+2 TCA intermediates, while entry via PC leads to M+3 TCA intermediates (Figure 4B). [U-^13^C]-glutamine is incorporated into glutamate and α-ketoglutarate (α-KG) as M+5, and α-KG oxidation leads to M+4 TCA labeling and reductive carboxylation results in M+3 TCA metabolites except for citrate, which is M+5 (Figure 4D). When we compared relative glucose incorporation into TCA intermediates via oxidative and reductive pathways, we found a small but significant decrease in glucose incorporation in NF1-45 cells (Figure 4C). Conversely, when we compared the relative glutamine incorporation into TCA intermediates via oxidative and reductive pathways, we saw a small but significant increase in glutamine incorporation in the NF1-45 cells (Figure 4E). This data suggests a small but statistically significant *NF1*-driven shift in TCA carbon source from glucose to glutamine. We next analyzed the total aqueous and organic metabolite pools and determined that *NF1* deficiency increased select TCA and REDOX intermediates and increased levels of many lipid species, particularly triglycerides (TG) (Figure 4F). This *NF1*- dependent increase in intracellular TG is consistent with previous findings in *NF1*-deficient muscle cells [40]. Next, we wanted to understand the impact of *NF1* deficiency on lipid class pools, and we found an increase in lipid levels across all classes except phosphatidylglycerol (PG), phosphatidylinositol (PI), phosphatidylserine (PS), and sphingomyelin (SM) (Figure 4G).

**Figure 4:**
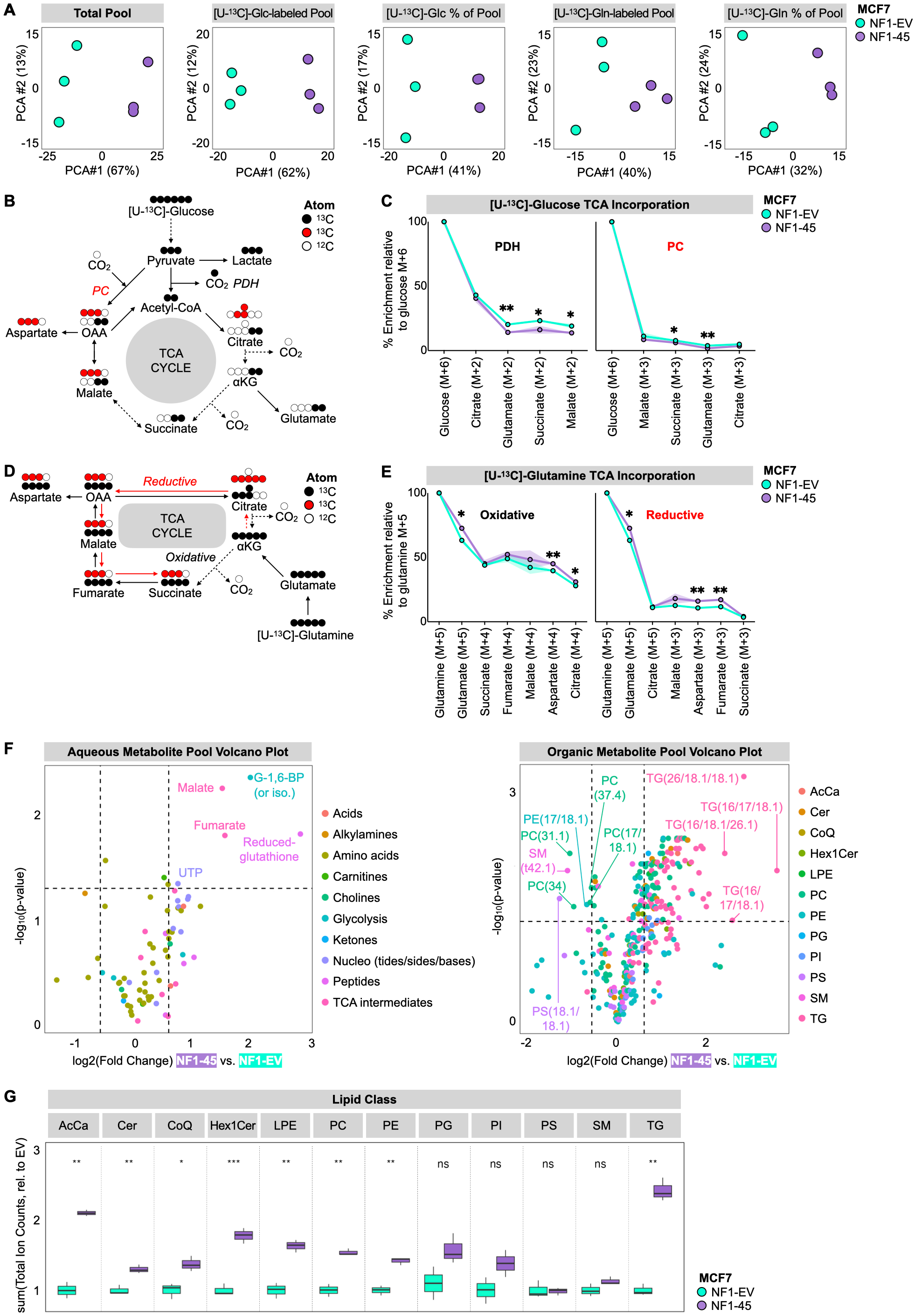
*NF1* deficiency increases intracellular lipid pools. A) *NF1* deficiency alters the ER+ breast cancer metabolome. PCA analysis of NF1-EV and *NF1*-deficient NF1-45 MCF7 LC-MS data (n=1, 3 technical replicates per biological replicate). B) Schematic of carbon flow from [U-13C]-glucose. C) *NF1* deficiency decreases glucose incorporation into TCA intermediates. Relative metabolite enrichment compared to M+6 [U-13C]-glucose (n=1, 3 technical replicates per biological replicate). D) Schematic of carbon flow from [U-13C]-glutamine. E) *NF1* deficiency increases glutamine incorporation into TCA intermediates. Relative metabolite enrichment compared to M+5 [U-13C]-glutamine (n=1, 3 technical replicates per biological replicate). F) *NF1* deficiency increases TCA and REDOX intermediates and intracellular lipid species. Volcano plots describing aqueous and organic metabolite pool sizes between NF1-45 and NF1-EV control (n=1, 3 technical replicates per biological replicate). The shape represents the geometric mean of the three technical replicates. G) *NF1* deficiency increases total intracellular lipids. Boxplot of the sum of ion counts normalized to the mean ion count of the NF1-EV control within each lipid class. The line represents the geometric mean of the three technical replicates and the box boundaries at the 1^st^ and 3^rd^ quartile boundaries (n=1, 3 technical replicates per biological replicate).

### 3.5 *NF1* mutation increases synergy between RAS pathway and triglyceride synthesis inhibitors

We next wanted to understand if metabolic inhibitors targeting NF1-driven metabolic changes synergize with traditional targeted inhibitors in *NF1*-deficient ER+ breast cancer cells. For targeted RAS and PI3K/AKT inhibitors, we evaluated the antiproliferative effects of Tamoxifen (4OHT), Cobimetinib (COB), and Everolimus (EVE) (Figure 1E; Supplementary Fig. 3A). For metabolic inhibitors, we evaluated the antiproliferative effects of the following metabolic inhibitors: 2DG, bis-2-(5- phenylacetamido-1,3,4-thiadiazol-2-yl)ethyl sulfide (BPTES), Etomoxir (EX), PF-04620110 (DGAT1i), and PF-06424439 (DGAT2i) (Figure 5A). We tested the glycolysis inhibitor, 2DG, because we saw increased glycolysis in NF1-45 cells (Figure 3D). 2DG inhibits glycolysis via competitive hexokinase inhibition, so we anticipated increased 2DG sensitivity in NF1-45 cells. We tested the glutaminase (GLS) inhibitor, BPTES, because we saw increased glutamine incorporation into TCA intermediates in NF1-45 cells (Figure 4E). GLS catalyzes the conversion of glutamine to glutamate, and glutamate dehydrogenase (GDH) catalyzes the conversion of glutamate to α-ketoglutarate, which can then enter the TCA cycle, so we anticipated increased BPTES sensitivity in NF1-45 cells. We next tested the carnitine palmitoyl transferase 1 (CPT1) inhibitor, Etomoxir (EX), because we saw increased palmitoyl and oleoyl carnitine (AcCa) species, intermediates in mitochondrial FA transport, in NF1-45 cells (Figure 4G). This increase in AcCa species suggested increased or impeded mitochondrial FA transport, a process facilitated by CPT1, so we anticipated increased Etomoxir sensitivity in NF1-45 cells. Lastly, we chose to test the diacylglycerol o-acyltransferase 1 and 2 (DGAT1, DGAT2) inhibitors, PF-04620110 (DGAT1i), and PF-06424439 (DGAT2i), because we saw a ∼2.5X increase in total TG species in NF1-45 cells (Figure 4G). DGATs catalyze the final step in TG synthesis, so we anticipated increased DGAT1i and DGAT2i sensitivity in NF1-45 cells. Because DGAT2 had ∼2X higher expression than DGAT1 in MCF7 cells, we chose to test DGAT2i as a single agent and DGAT1/2i in combination (Supplementary Fig. 3B). When comparing single-agent antiproliferative effects, we found that NF1-45 cells were more sensitive to ERα inhibition by 4OHT, mitogen-activated protein kinase kinase (MEK) inhibition by COB, and mechanistic target of rapamycin (mTOR) inhibition by EVE (Figure 5B). NF1-45 cells were also more sensitive to each metabolic inhibitor tested (Figure 5B). Next, we combined each targeted and metabolic inhibitor to investigate genotype-driven synergy changes using a Bliss Synergy model and the SynergyFinder R package (Figure 5 and Supplementary Fig. 3C-G). A synergy score of less than -10 indicates that the drug-drug interaction is likely antagonistic, a synergy score between -10 and 10 indicates that the drug-drug interaction is likely additive, and a synergy score greater than 10 indicates that the drug-drug interaction is likely synergistic [55]. When we evaluated 2DG synergy with RAS, ER, and PI3K/AKT inhibitors, we saw a ∼1.5X *NF1*-dependent increase in synergy between 2DG and 4OHT, COB, and EVE (Figure 5C, Supplementary Fig. 3C). This *NF1*-dependent increase in 2DG- targeted inhibitor synergy could be due to increased glycolytic glucose use, but the loss of lipid, nucleotide, and TCA precursor species could also account for *NF1*-driven synergy. When we evaluated BTPES synergy with RAS, ER, and PI3K/AKT inhibitors, we saw ∼2X more synergy between BPTES and 4OHT and ∼3.5X more synergy between BPTES and COB in the *NF1*-deficient NF1-45 cells (Figure 5D, Supplementary Fig. 3D). When we evaluated Etomoxir synergy with RAS, ER, and PI3K/AKT inhibitors, we saw ∼1.5X more synergy Etomoxir and 4OHT, ∼2X more synergy between Etomoxir and COB, and a small increase in synergy between Etomoxir and EVE in NF1-45 cells (Figure 5E, Supplementary Fig. 3E). When we evaluated DGAT2i synergy with RAS, ER, and PI3K/AKT inhibitors, we saw a ∼1.5-2X increase in synergy between DGAT2i and COB and a ∼3X increase in synergy between DGAT2i and EVE in NF1-45 cells (Figure 5F, Supplementary Fig. 3F). When we evaluated DGAT1/2i synergy with RAS, ER, and PI3K/AKT inhibitors, we saw a ∼4X increase in synergy between DGAT1/2i and 4OHT, a ∼1.5X increase in synergy between DGAT1/2i and COB, and a 1.5X increase in synergy between DGAT1/2i and EVE (Figure 5G, Supplementary Fig. 3G). Taken together, these results underscore the *NF1*-driven energetic and metabolic changes seen in our Seahorse XF-96 and mass spectrometry experiments. We also provide evidence connecting *NF1*- driven metabolic reprogramming to inhibitor sensitivity and synergy. Indeed, *NF1*-deficient NF1-45 cells have increased glycolysis and increased glycolysis inhibitor sensitivity and synergy, increased TCA glutamine incorporation and increased GLS inhibitor sensitivity and synergy, increased AcCa pools and increased CPT1 inhibitor sensitivity and synergy, and increased TG pools and increased DGAT1/2 inhibitor sensitivity and synergy. When we compared the antiproliferative effects of all our combination strategies (Supplementary Fig. 4-5) and their respective Bliss synergies (Figure 5 and Supplementary Fig. 3C-G), we found that inhibition of RAS and triglyceride synthesis most effectively halted proliferation in *NF1*-deficient NF1-45 cells (Figure 5H). These findings pave the way for future *in vivo* experiments and highlight potential combination treatments for *NF1*-deficient, ER+ breast cancer.

**Figure 5:**
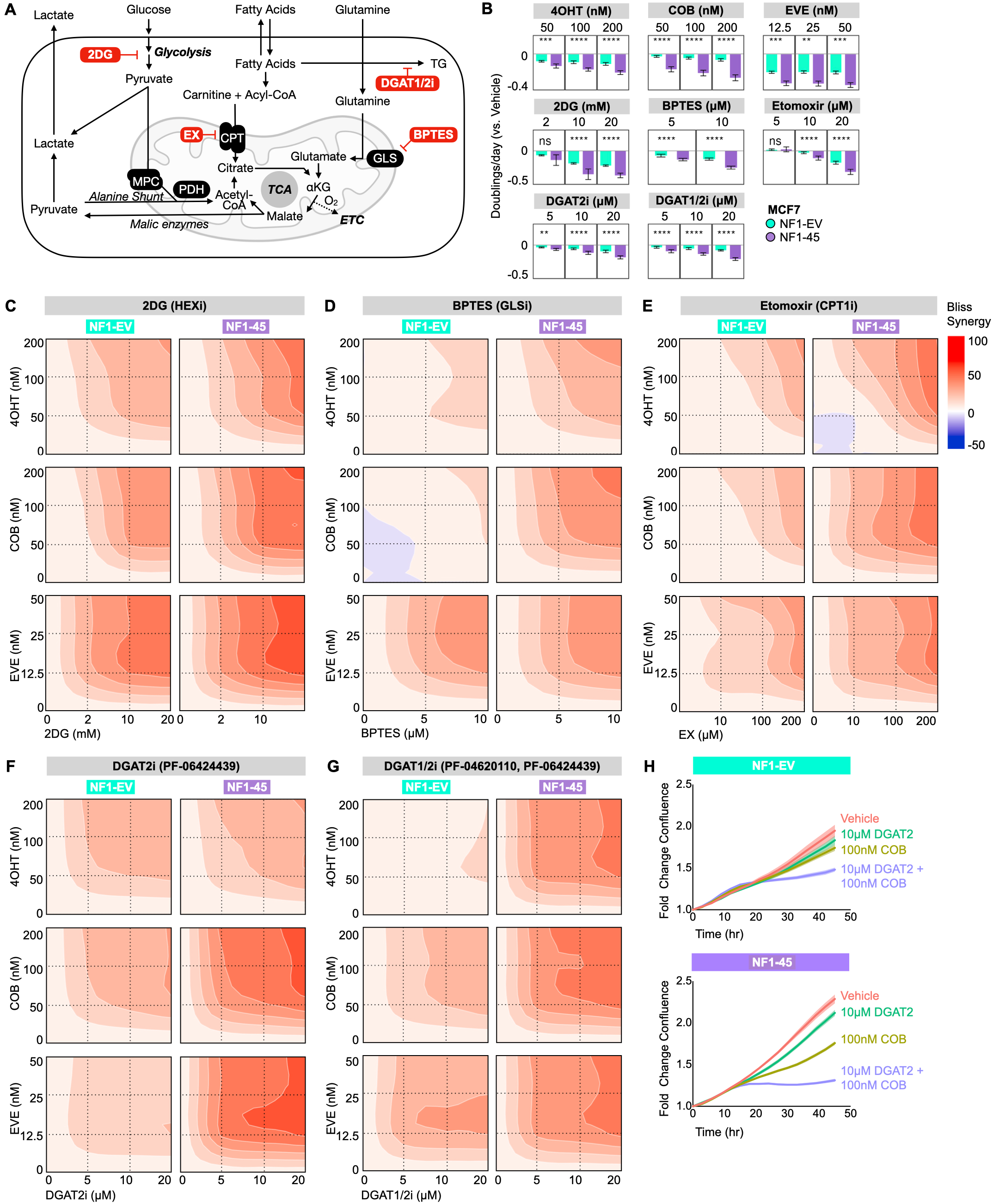
*NF1* mutation increases synergy between RAS pathway and triglyceride synthesis inhibitors. A) Schematic description of selected metabolic inhibitors and their respective enzymatic targets: 2-Deoxy-D-glucose (2DG), bis-2-(5-phenylacetamido-1,3,4-thiadiazol-2-yl)ethyl sulfide (BPTES), Etomoxir (EX), PF-04620110 (DGAT1i), PF-06424439 (DGAT2i). B) *NF1* deficiency increases sensitivity to targeted and metabolic inhibitors. A crossbar plot describing the antiproliferative impact of 4OHT, COB, EVE, 2DG, BPTES, EX, DGAT2i, and DGAT1/2i on NF1-EV and NF1-45 (n=2, 8 technical replicates per biological replicate). The bar represents the geometric mean, and the error bar boundaries represent the 95% confidence interval boundaries. C) Bliss synergy plots describing the synergy between 2DG and 4OHT, COB, and EVE. Bliss synergy was calculated and visualized using the SynergyFinder R package (n=2, 8 technical replicates per biological replicate). D) Bliss synergy plots describing the synergy between BPTES and 4OHT, COB, and EVE. Bliss synergy was calculated and visualized using the SynergyFinder R package (n=2, 8 technical replicates per biological replicate). E) Bliss synergy plots describing the synergy between Etomoxir and 4OHT, COB, and EVE. Bliss synergy was calculated and visualized using the SynergyFinder R package (n=2, 8 technical replicates per biological replicate). F) Bliss synergy plots describing the synergy between DGAT2i and 4OHT, COB, and EVE. Bliss synergy was calculated and visualized using the SynergyFinder R package (n=2, 8 technical replicates per biological replicate). G) Bliss synergy plots describing the synergy between DGAT1i and DGAT2i combination treatment and 4OHT, COB, and EVE. Bliss synergy was calculated and visualized using the SynergyFinder R package (n=2, 8 technical replicates per biological replicate). H) IncuCyte proliferation graphs of EV and MUT45 cells grown in 10nM E2 and Vehicle, 10μM DGAT2, 100nM COB, or 10μM DGAT2 + 100nM COB. The geometric mean is visualized by the connecting line and the 95% confidence interval by the shading (n=2, 8 technical replicates per biological replicate).

## DISCUSSION

Historically, the tumor suppressor *NF1* was studied in the context of NF-associated benign and malignant neuronal tumors, but recent genomic and epidemiological analyses have identified *NF1* as a driver of inherited breast cancer, sporadic breast cancer, and endocrine resistance [2–10]. Despite these recent analyses connecting *NF1* and breast cancer, there is a fundamental need for a greater understanding of neurofibromin’s role in breast cancer development, biology, and therapeutic response. One essential and previously unexplored area of study is the relationship between *NF1* deficiency and breast cancer cell metabolism. Activated oncogenes and inactivated tumor suppressors are known to contribute to tumorigenesis by reprogramming cellular energetics and metabolism towards macromolecular synthesis – a phenomenon termed “metabolic reprogramming” [21]. Despite neurofibromin’s known role as a breast cancer tumor suppressor, no study had evaluated *NF1*-driven metabolic reprogramming in ER+ breast cancer – a gap this study aimed to fill.

We employed a novel *NF1*-deficient ER+ breast cancer MCF7 model and previously published *Nf1*-deficient rat models to evaluate *NF1*-driven metabolic reprogramming in ER+ breast cancer. We demonstrated that *NF1* deficiency results in decreased *NF1* mRNA and decreased neurofibromin 250kDa isoform expression. This change in neurofibromin expression is accompanied by a decrease in estrogen pathway signaling and an increase in RAS and PI3K/AKT pathway signaling, which is expected given neurofibromin’s RAS-GAP function. We also demonstrate that *NF1* deficiency increases 2D and 3D cell proliferation while reducing basal and maximal oxidative ATP production twofold.

Using RNAseq, we found that *NF1* deficiency globally altered transcription. When we investigated the impact of *NF1* deficiency on metabolism-specific transcription, we noted a change in OXPHOS-related transcription and FA and lipid-related transcription. When we performed glucose and glutamine stable isotope labeling, we saw a decrease in glucose TCA incorporation and an increase in glutamine TCA incorporation, suggesting a shift in TCA intermediate carbon sourcing. Perhaps the most striking metabolic finding was a ∼1.5-2.5-fold increase in lipid species across classes. Of the enriched species, TGs were most dramatically and consistently increased.

Lastly, we demonstrated that *NF1*-driven metabolic changes introduce therapeutic vulnerabilities and drug synergies. We illustrated increased sensitivity to single-agent inhibitors against ER, RAS, and PI3K/AKT pathway members and increased sensitivity to glycolysis, glutamine, fatty acid, and lipid metabolism inhibitors that parallel *NF1*-driven pathway and metabolic changes. When we combined targeted and metabolic inhibitors, we saw broad *NF1*-driven synergy between glutamine, fatty acid mitochondrial transport, and triglyceride synthesis inhibitors and ER, RAS, and PI3K/AKT pathway inhibitors. When we evaluated our drug combinations for both antiproliferative *and* synergistic effects, we determined that RAS and triglyceride synthesis inhibition was the most effective strategy to inhibit proliferation in *NF1*-deficient breast cancer cells. This study is the first to combine RAS and triglyceride synthesis inhibitors in an RAS-active tumor model, let alone an *NF1*-deficient, ER+, RAS-active context.

We found a significant phenotypic overlap between *NF1*-deficiency and RAS-activation-driven metabolic reprogramming. In both of our *NF1*-deficient systems and published RAS-active systems, we see increased cellular proliferation, increased glutamine TCA incorporation, and increased neutral lipid accumulation (TG). We also saw increased glutaminase and CPT1 inhibitor sensitivity in our *NF1*- deficient models and published *Nf1*-deficient and RAS-active models [43; 56]. We have demonstrated that *NF1*-deficient ER+ breast cancer cells have impaired mitochondrial respiration without a change in total mitochondrial load, an observation reproduced in *Nf1^-/-^* mouse embryonic fibroblasts and *Nf1-KO drosophila melanogaster* [36; 37]. Published data from RAS-active cells shows an overlapping energetic phenotype with impaired mitochondrial respiration but increased mitochondrial load. This discrepancy in mitochondrial load may speak to a difference in the degree of RAS pathway activation, specifically to the difference between an *NF1*-deficient RAS-dysregulated system and constitutively RAS-activated systems. The difference in mitochondrial load could also relate to the fundamental signaling distinction between an *NF1*-deficient and a RAS-active system: the impact of *NF1* deficiency on ERα at the transcriptional activity. Neurofibromin is a known ERα transcriptional co-repressor, so *NF1* deficiency can impact ER’s transcriptional activity. In our *NF1*-deficient NF1-45 cells, we see decreased ERK-dependent S118 phosphorylation despite increased ERK T202/Y204 phosphorylation and decreased total ERα, highlighting the nuanced relationship between *NF1* deficiency and estrogen signaling. Overall, want to underscore the importance of further metabolic inquiry in *NF1*-deficient contexts given the efficacy of metabolism-directed therapeutics in RAS-active cancers.

## CONCLUSION

In conclusion, this study demonstrates that *NF1* deficiency drives metabolic reprogramming in ER+ breast cancer. *NF1*-driven ER+ breast cancer metabolic reprogramming is characterized by increased proliferation, constrained oxidative ATP production, increased glutamine TCA influx, and lipid pool expansion. These metabolic changes introduce novel single-agent sensitivity and metabolic-to-targeted inhibitor synergies. This study highlights neurofibromin’s ability to alter cellular metabolism across cell types and model organisms.

## List of Abbreviations

2DG: 2-Deoxy-D-glucose
4OHT: 4-Hydroxytamoxifen or tamoxifen
ATP: Adenosine Triphosphate
BPTES: Bis-2-(5-phenylacetamido-1,3,4-thiadiazol-2-yl)ethyl sulfide
COB: Cobimetinib
CPT1: carnitine palmitoyl transferase 1
CRISPR: Clustered Regularly Interspaced Short Palindromic Repeats
DGAT1: Diacylglycerol O-Acyltransferase 1
DGAT1/2i: Diacylglycerol O-Acyltransferase 1 and 2 Inhibitor
DGAT2: Diacylglycerol O-Acyltransferase 2
DMEM: Dulbecco’s Modified Eagle Medium
dFBS: Dialyzed Fetal Bovine Serum
ECAR: Extracellular Acidification Rate
EMT: Epithelial-Mesenchymal Transition
ER+: Estrogen Receptor Positive
ETC: Electron Transport Chain
EVE: Everolimus
EX: Etomoxir
FA: Fatty acid
FBS: Fetal bovine serum
GAP: GTPase activating protein
GDH: Glutamate dehydrogenase
GLS: Glutaminase
GSEA: Gene Set Enrichment Analysis
GRD: GAP-related domain
H-ESI: Heated Electrospray Ionization
KEGG: Kyoto Encyclopedia of Genes and Genomes
KO: Knock-out
LC-MS: Liquid Chromatography-Mass Spectrometry
mRNA: Messenger RNA
mTOR: Mechanistic Target of Rapamycin
MEK: Mitogen-Activated Protein Kinase Kinase
MPC: mitochondrial pyruvate carrier
*Nf1*: Neurofibromin 1 (rat) gene
NF: Neurofibromatosis Type 1
*NF1*: Neurofibromin 1 (human) gene
OCR: Oxygen Consumption Rate
OXPHOS: Oxidative Phosphorylation
PCA: Principal Component Analysis
PCR: Polymerase Chain Reaction
PG: Phosphatidylglycerol
PI: Phosphatidylinositol
PS: Phosphatidylserine
PI3K-AKT: Phosphoinositide 3-kinase-AKT pathway
RAS: Rat sarcoma viral oncogene homolog
REACTOME: REACTOME pathway database
RNAseq: RNA Sequencing
SM: Sphingomyelin
TCA: Tricarboxylic Acid Cycle
TG: Triglyceride
VDAC: Voltage-dependent Anion Channel

## AUTHOR CONTRIBUTIONS

RH wrote the manuscript. CG, MS, and EL provided edits for the manuscript. RH performed all experiments except the following: ET performed the ER, RAS, and PI3K signaling Western Blot analysis and the RNAseq extraction, AE ran the LCMS and MS machines. RH analyzed the data for all experiments. AE and RS provided mass spectrometry analysis consultation. IB performed RNAseq preprocessing. RS and EL contributed analytical tools. PD designed the MCF7 CRISPR guides. CG and MS obtained funding. All authors have given final approval of the final manuscript version for publication.

## ACKNOWLEDGEMENTS

We would like to thank the VAI Vivarium staff, Flow Cytometry Core, and Genomics Core for their valued expertise. We want to extend our deepest gratitude to all VAI Mass Spectrometry Core members for their expertise, camaraderie, and mentorship. We also want to thank Rae’s thesis advisory committee members, Russel (Rusty) Jones, PhD, Ralph DeBerardinis, PhD, and Timothy Triche, PhD, for their direction and counsel throughout this project. Lastly, we would like to thank the VAI Department of Metabolism and Nutritional Programming for their metabolic support, especially the Lien Lab.

