## Supplementary Figures for "*NF1* deficiency drives metabolic reprogramming in ER+ breast cancer"

Supp. Figure 1

**A iNFiXion BioScience neurofibromin iNF-07E antibody epitope schematic**

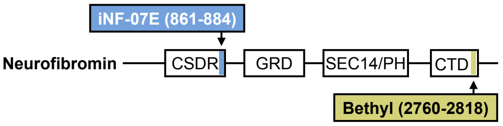


**B Neurofibromin iNF-07E antibody IHC signal is *NF1*-dependent**

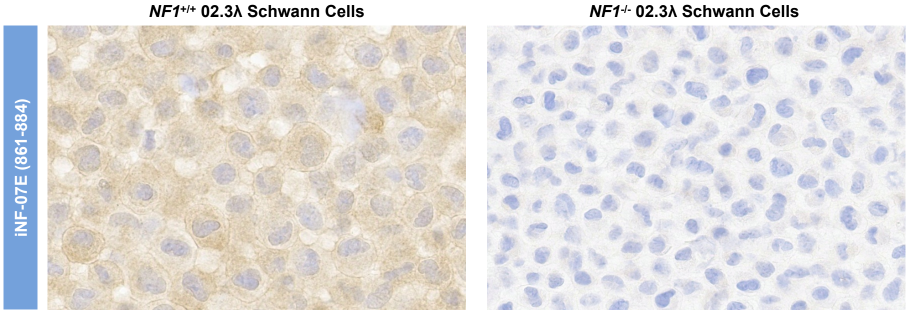


**C Neurofibromin iNF-07E antibody is specific to neurofibromin and preferentially binds neurofibromin at 250 kDa**

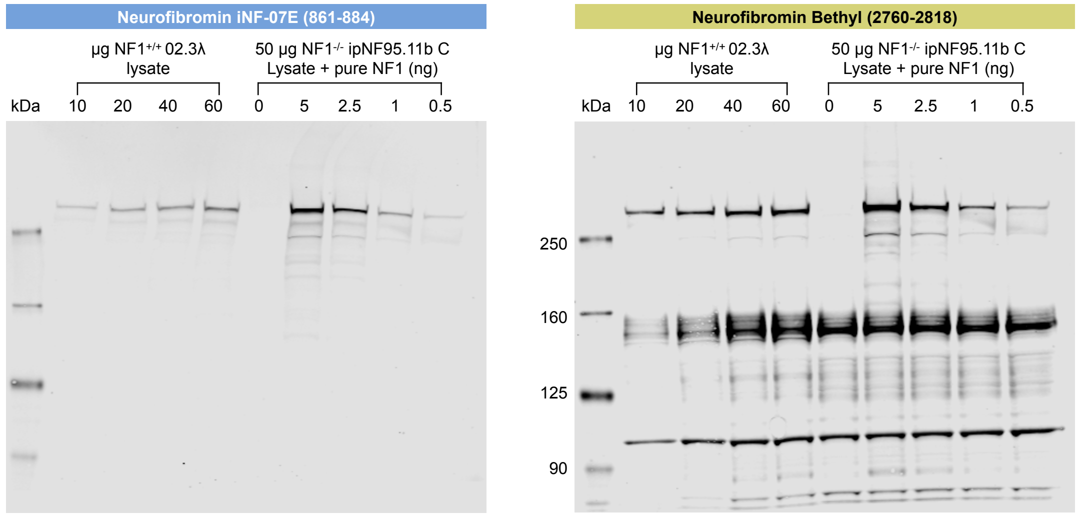


**Supplementary Figure 1: iNFiXion NF1 iNF-07E antibody data**
A) iNFiXion BioScience neurofibromin iNF-07E antibody epitope schematic. The iNF-07E neurofibromin antibody binds at amino acids 861-884 compared to the Bethyl neurofibromin antibody, which binds at amino acids 2760-2818. Data provided by iNFiXion BioScience. B) iNFiXion BioScience neurofibromin iNF-07E antibody IHC signal is *NF1*-dependent. Data provided by iNFiXion BioScience. C) iNFiXion BioScience neurofibromin iNF-07E antibody western blot signal is specific to neurofibromin and preferentially binds neurofibromin at 250kDa. Data provided by iNFiXion BioScience.

**Supp. Figure 2**

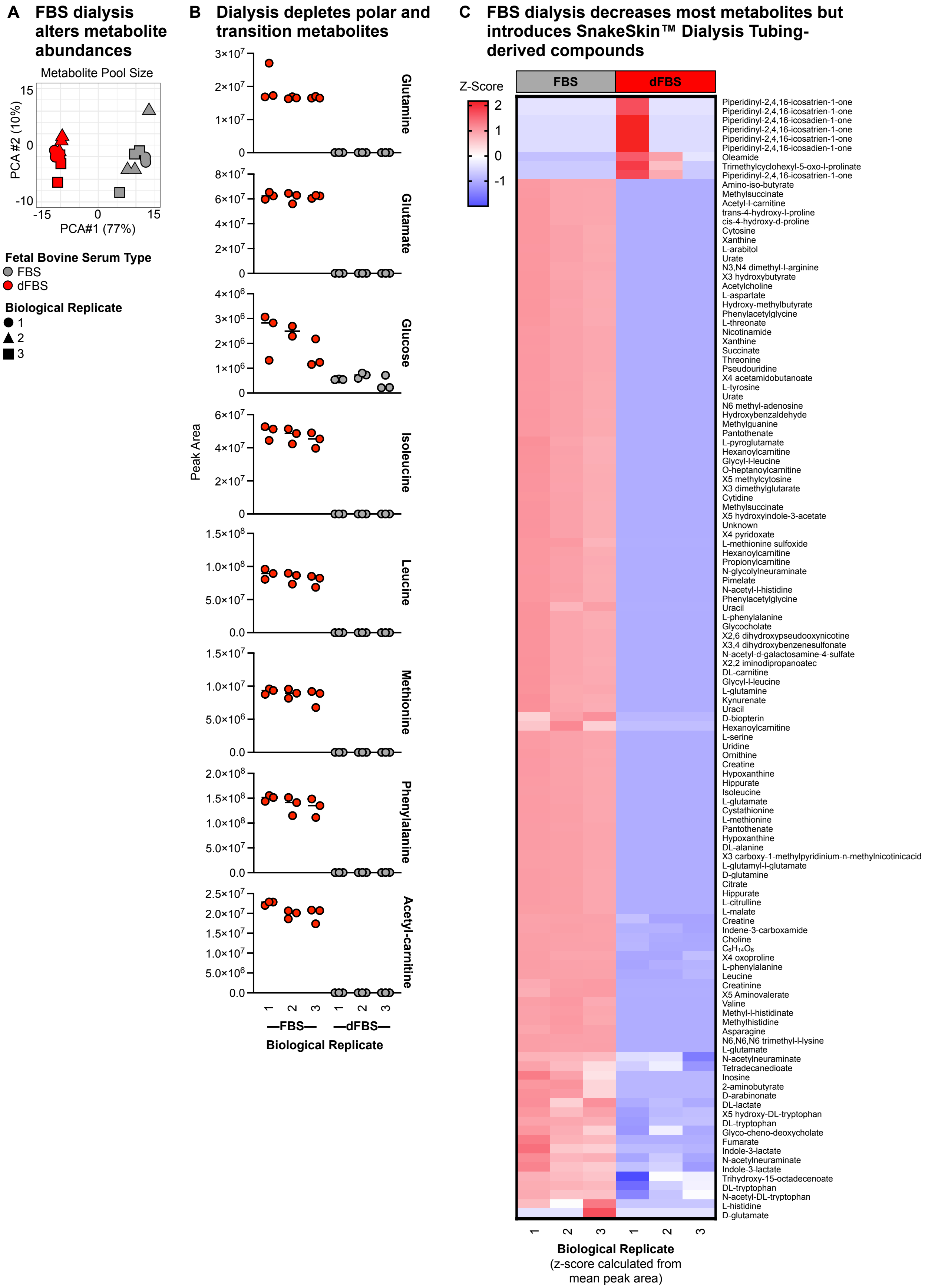


**Supplementary Figure 2: FBS dialysis depletes most measured metabolites**
A) PCA analysis complete vs. dialyzed FBS (n=3, 3 technical replicates per biological replicate). B) Comparison of selected metabolite peak values between complete and dialyzed FBS (n=3, 3 technical replicates per biological replicate). C) Comparison of relative abundance of all measured and identified metabolites between complete and dialyzed FBS (n=3, 3 technical replicates per biological replicate).

### A MCF7 Inhibitor Dosage References

**Supp. Figure 3**

| Drug | Dose Range | Publication | Metabolic Effect | Proliferative / Viability Effect |
|---|---|---|---|---|
| Tamoxifen | 100nM | Yee et al. 2017 | NA | ~50% reduction in cell viability |
| Cobimetinib | 100nM | Mills et al. 2023 | NA | ~25% reduction in cell viability |
| Everolimus | 25nM | Lewis-Wambi et al. 2016 | NA | 50% reduction in cell viability |
| 2DG | 5mM | Prehn et al. | Increased intracellular glucose | NA |
| BPTES | 10µM | Jeong et al. 2019 | No change in intracellular glutamine, decreased GLS protein expression | No proliferative or viability effect |
| | 10µM | Di et al. 2015 | Decreased glutamine uptake, decreased ATP | NA |
| | 20µM | Raftery et al. 2018 | Decreased intracellular glutamate | NA |
| Etomoxir | 10-200µM | Patti et al. 2018 | Inhibited fatty acid oxidation at 10µM, Complex I inhibition at 200µM | Decreased cell proliferation at 200µM |
| PF-06424439 | 10µM | Frasor et al. 2018 | Decreased triglycerides after treatment (combo DGAT1/2i) | Decreased final confluency |
| PF-04620110 | 10µM | Seco et al. 2021 | Decreased lipid droplets, altered lipid gene expression | Decreased cell proliferation at 100µM |
| | 10µM | Frasor et al. 2018 | Decreased triglycerides after treatment (combo DGAT1/2i) | Decreased final confluency |

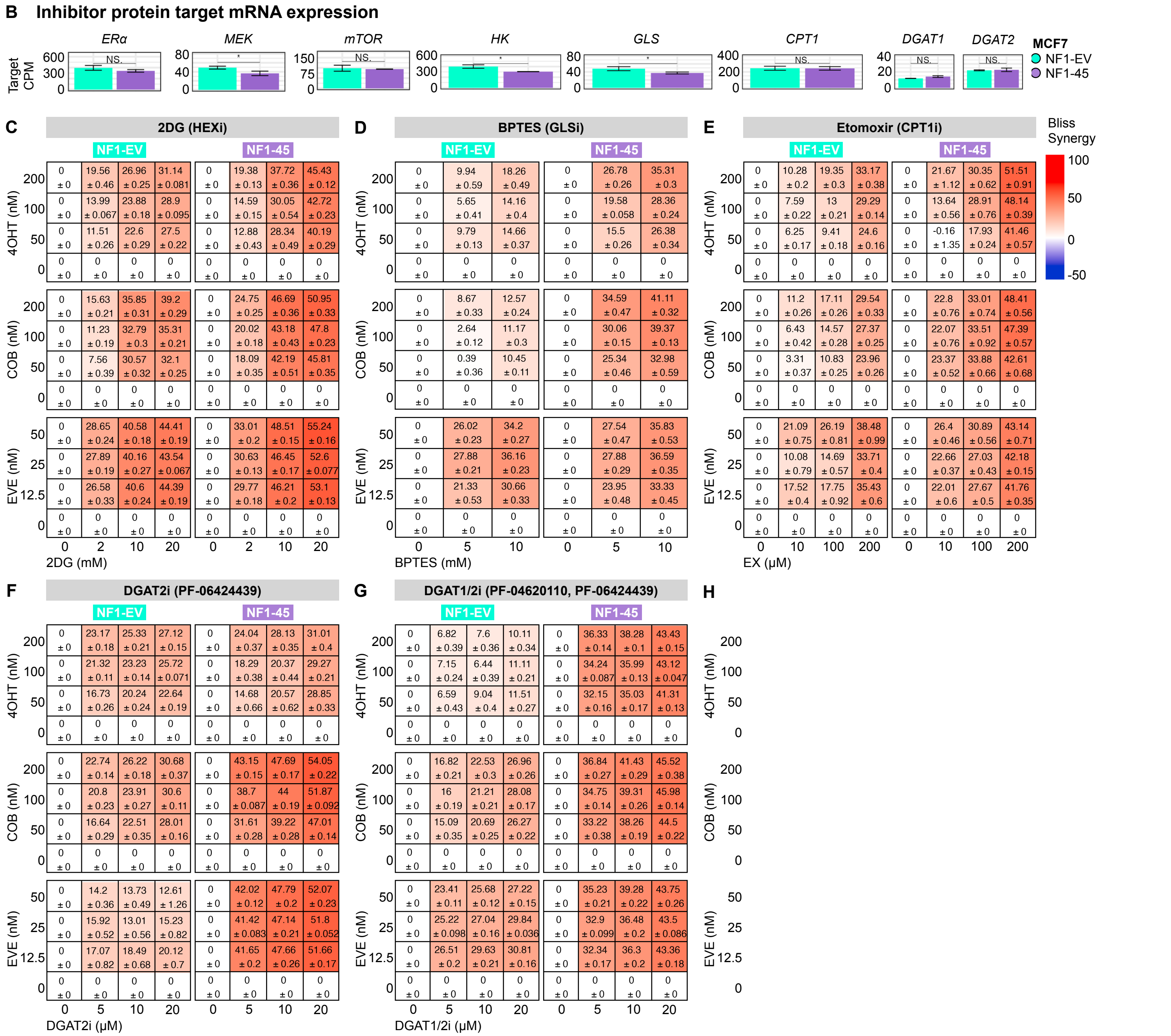


**Supplementary Figure 3: Drug synergy data**

A) MCF7-specific targeted and metabolic inhibitor dose reference table. B) Mean inhibitor target mRNA expression (n=3) The bar represents the geometric mean, and the error bar boundaries represent the standard deviation. C) Raw Bliss synergy plots describing the synergy between 2DG and 4OHT, COB, and EVE. Bliss synergy was calculated and visualized using the SynergyFinder R package (n=2, 8 technical replicates per biological replicate). D) Raw Bliss synergy plots describing the synergy between BPTES and 4OHT, COB, and EVE. Bliss synergy was calculated and visualized using the SynergyFinder R package (n=2, 8 technical replicates per biological replicate). E) Raw Bliss synergy plots describing the synergy between EX and 4OHT, COB, and EVE. Bliss synergy was calculated and visualized using the SynergyFinder R package (n=2, 8 technical replicates per biological replicate). F) Raw Bliss synergy plots describing the synergy between DGAT2i and 4OHT, COB, and EVE. Bliss synergy was calculated and visualized using the SynergyFinder R package (n=2, 8 technical replicates per biological replicate). G) Raw Bliss synergy plots describing the synergy between DGAT1i and DGAT2i combination treatment and 4OHT, COB, and EVE. Bliss synergy was calculated and visualized using the SynergyFinder R package (n=2, 8 technical replicates per biological replicate).

**A** Raw doublings per day values for Bliss Synergy analysis

**Supp. Figure 4**

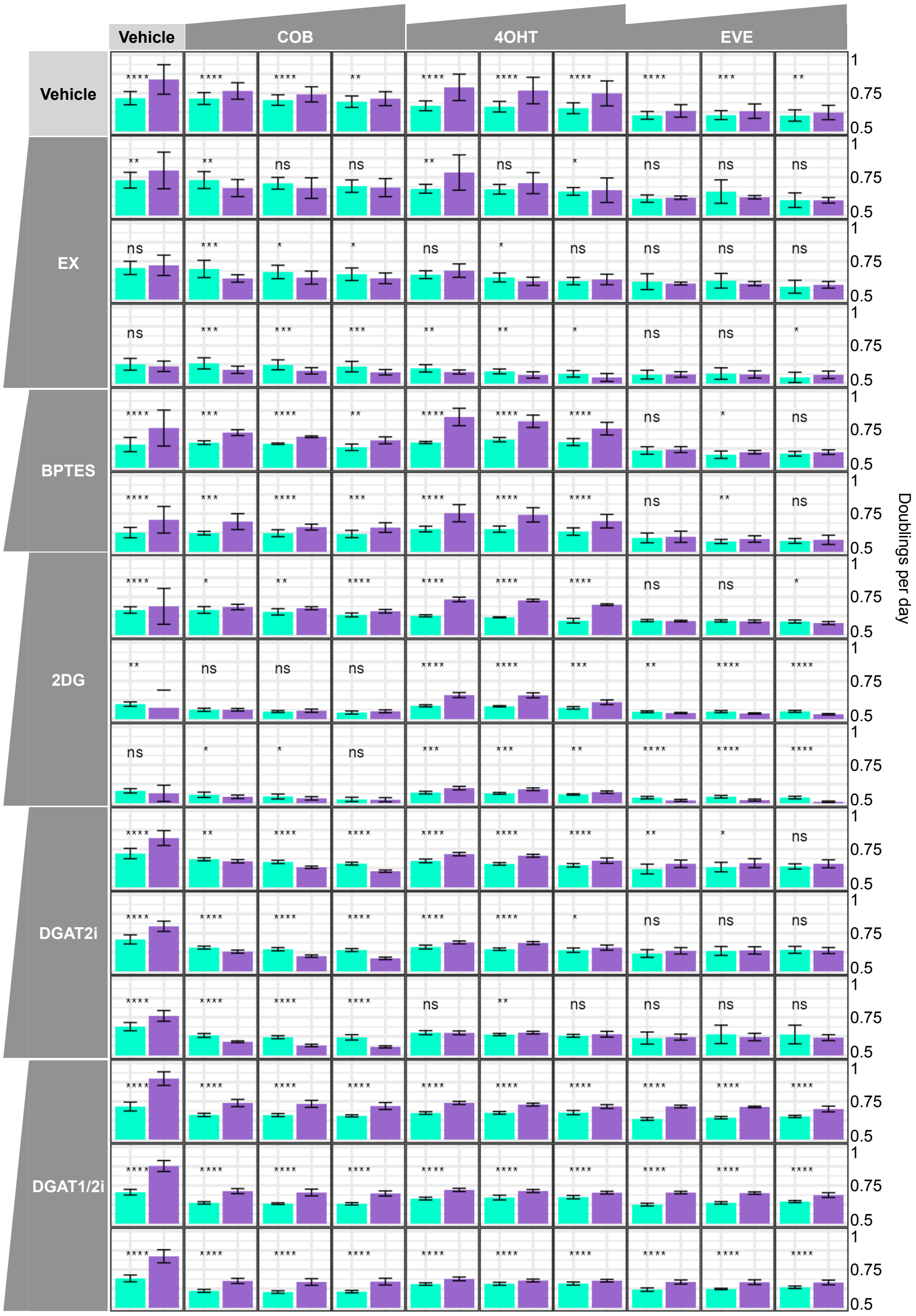


**Supplementary Figure 4: Raw drug synergy doublings per day data**

A) Raw doublings per day values for combination inhibitor treatments. The bar represents the geometric mean, and the error bar boundaries represent the standard deviation (n=2, 8 technical replicates per biological replicate).

A Normalized doublings per day values for Bliss Synergy analysis

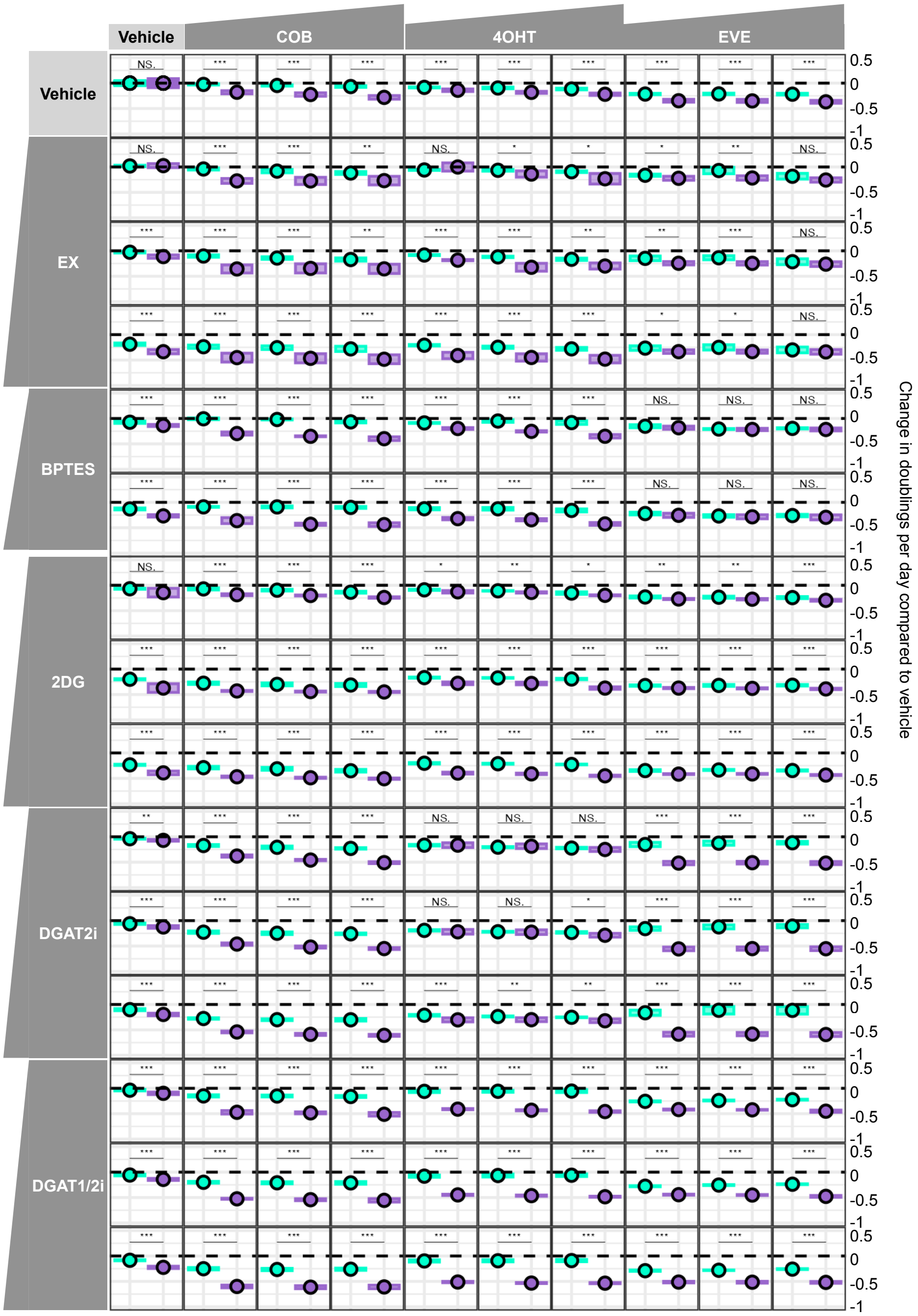


**Supplementary Figure 5: Normalized doublings per day values for Bliss Synergy analysis**
A) Normalized doublings per day values for combination inhibitor treatments used for Bliss synergy analysis. The symbol represents the geometric mean, and the crossbar boundaries represent the 95% confidence interval (n=2, 8 technical replicates per biological replicate).
